# Analyzing Drumming Technique with Trajectory Optimization

**DOI:** 10.1101/2023.10.09.561270

**Authors:** Owen Douglas Pearl

## Abstract

Percussion began as a global phenomenon over seven thousand years ago and has continued to develop and shape human culture to this day. While drumming techniques have been qualitatively optimized in highly competitive environments like international orchestral, jazz, and marching arts competitions, little quantitative optimization has been performed to maximize technique efficiency and few tools currently exist to make a thorough quantitative analysis possible. Here, I demonstrate how trajectory optimization can be applied to the domain of percussion to (1) identify areas of suboptimality in experimental drumming strokes, (2) search for alternative locally optimal techniques, (3) analyze the sensitivity of optimal drumming techniques to variation in a drummer’s body type, and (4) analyze trends across different stroke types to create generalizable drumming strategies for a more coherent and efficient approach to drumming. Each of these quantifiable outcomes is interpreted to provide teachable insights for percussionists that are difficult to distill simply using the human eye and qualitative feedback. I also provide an open-source codebase for efficiently performing trajectory optimization on a biomechanical drumming arm model so that others can adapt this methodology for further biomechanical analysis and pedagogical development.

## 1. Introduction

Percussion is a highly dynamic, physically demanding, and often precisely synchronized artform that began influencing human society as early as 5000 BC and has since developed into many rich forms of artistic expression, performance, and competition^[1]^. Some forms of drumming competition, such as those taking place in the context of the marching arts, are so dynamic and physically demanding that attempts to measure energy expenditure during a performance yield results similar to that of a trained marathon runner halfway through a marathon^[2]^. Because of the extremely selective nature of competitive drumming environments, from premier drumlines to world class orchestras and famous drum set players, drumming techniques and percussion instrument design have been artistically refined to produce the highest quality sound for nearly every musical context. However, given the complexities of the dynamic system of the arm, an artistic and qualitative approach to characterizing the best drumming technique is likely insufficient for fully understanding the advantages and disadvantages of selecting one technique or style over another. As a result, different percussion groups performing at comparable levels have adopted distinguished approaches to achieving the same basic goal of precisely controlling the motion of the bead of the drumstick to strike the drumhead at a desired time with a desired force^[3,4]^. While some quantitative analyzes of drumming systems have been performed either to better understand the system experimentally^[5,6]^ or to reproduce a controllable drumming motion with a robot^[7,8]^, the optimal drumming technique, or control policy, for driving the bead of a drumstick to a given state while minimizing the effort of the percussionist remains to be quantitatively investigated. Furthermore, the tools for completing such an investigation are not currently accessible to nonexperts of chaotic multibody dynamic systems and optimal control theory.

In this work, I formulate the problem of finding the minimum effort required to achieve a desired path of a drumstick as a trajectory optimization^[9]^ problem with a reduced-order four degree of freedom biomechanical drumming model and solve the problem with direct collocation (**Figure 1**) in order to illustrate how trajectory optimization techniques can be applied to the domain of percussion to facilitate teachable insights. First, I compare the optimal minimum effort solution from direct collocation against an experimental drum stroke performed by a drummer with 20 years of experience to evaluate whether trajectory optimization can be used to identify areas of suboptimality in the drummer’s technique. I hypothesize that the experimental drum stroke performed by the drummer will require more effort than the optimal technique. Next, I explore the impact of allowing flexibility on the tracking of the bead of the drumstick in the anterior-posterior direction to see if trajectory optimization can be used to identify alternative locally optimal techniques that might provide unique advantages. I hypothesize that tracking only the height of the bead of the drumstick will result in an optimal technique with a lower effort requirement than that which results from tracking the entire two-dimensional trajectory, Then, I analyze the differences between the optimal techniques for representative models of the average male and female body segment geometries and inertias to investigate the sensitivity of the optimal technique to body type. I hypothesize that the optimal technique identified using the body segments of the average female will require greater flexion at the elbow than the optimal technique using the body segments of the average male. Finally, I search for trends in the optimal technique across different strokes (single, double, and triple) to see if there are coherent patterns in joint function between different stroke types that can be used to inform real-world drumming strategy. I hypothesize that drumming trajectories for an increasing number of subsequent strokes will require more negative actuation at the shoulder and elbow.

**Figure 1.**
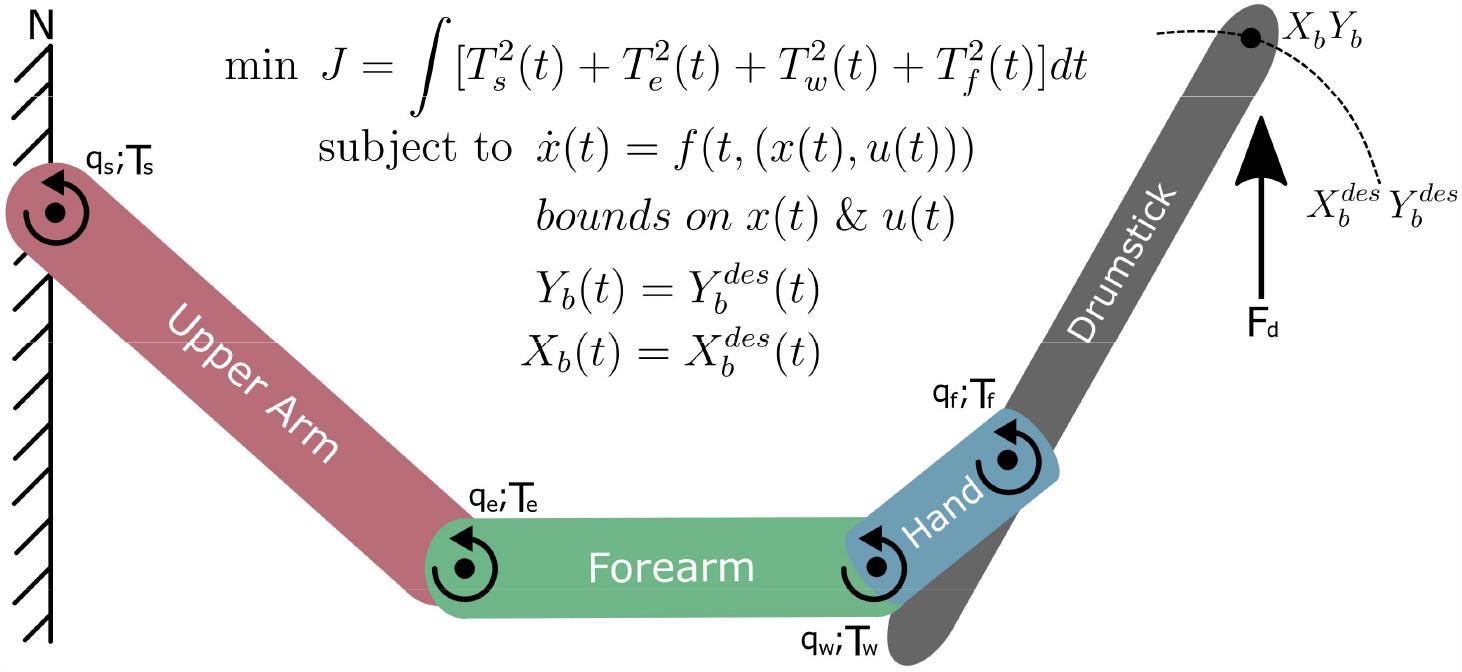
Trajectory Optimization with a Biomechanical Drumming Arm Model. The drumming arm model consists of four sagittal plane revolute joints and four idealized torque actuators, as well as one vertical reaction force applied at the bead of the drumstick. The solution to the trajectory optimization problem is the set of states (kinematics) and controls (dynamics) that minimizes the sum of the squared torque inputs required to track the desired two-dimensional trajectory of the bead of the drumstick subject to the nonlinear dynamics of the model and boundaries on the controls and states constraining them to the physiological limits of the human arm.

## 2. Materials and Methods

### 2.1 Collecting Experimental Drumming Data

Video of the author of this work (**Figure *2***) performing a downstroke at 100 beats per minute was recorded via an iPhone 8 Plus (Apple, Inc.; Cupertino, CA) at 240 frames per second (picture owned and approved by the author). The author is a 25-year-old male with 21 years of formal training spanning orchestral percussion, marching percussion, and jazz drum set, who most recently performed as a member of the Temple University Percussion Ensemble. The drum stroke was recorded by tracking the position of four retroreflective markers placed on the shoulder, elbow, wrist, and fulcrum joint centers as well as reflective tape wrapped on the bead of the drumstick.

**Figure 2.**
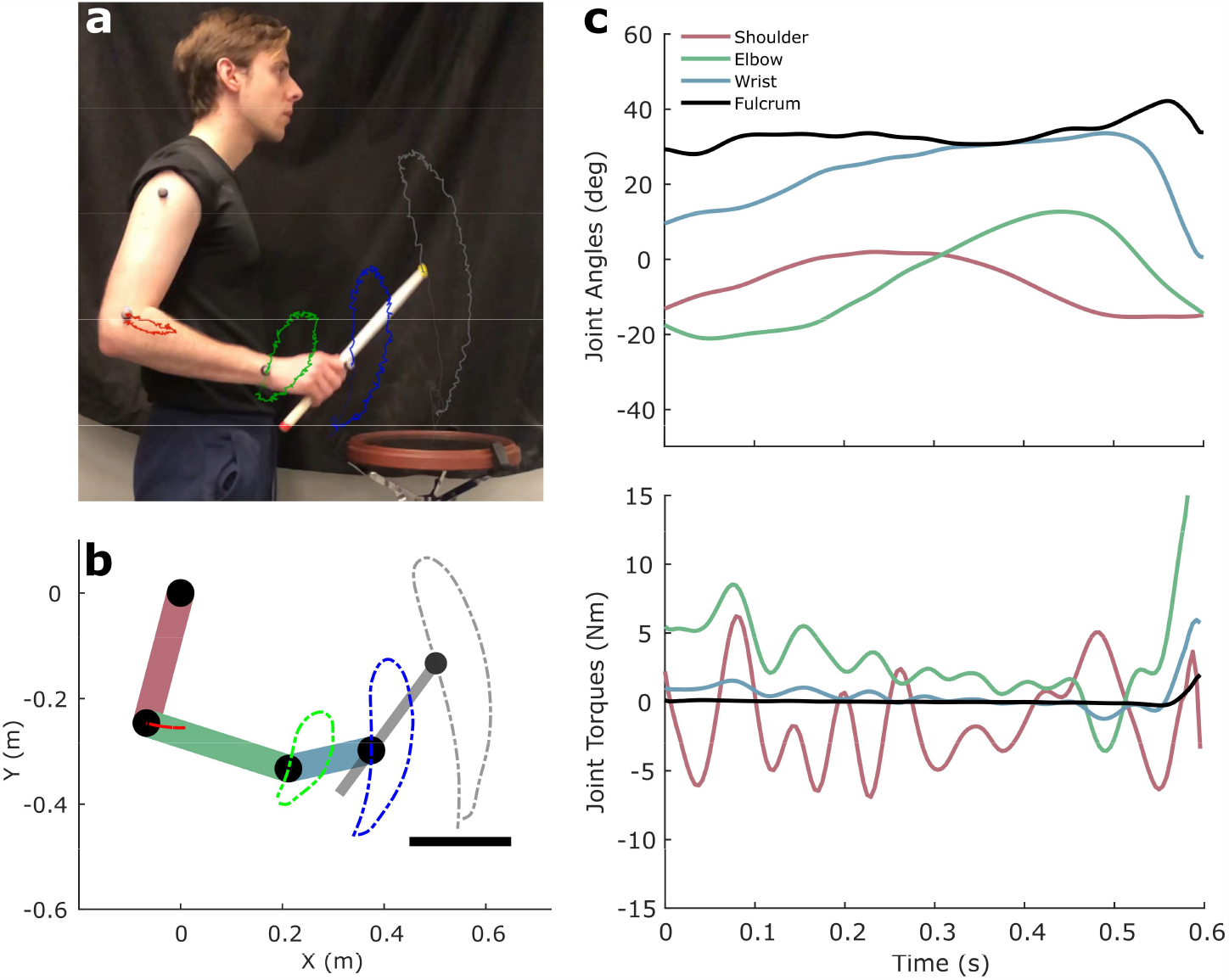
Biomechanical Analysis on Experimental Drumming Strokes. (**a**) Traces of the experimentally tracked markers placed on anatomical landmarks of the arm (picture of author owned and approved by the author) are shown during a downstroke performed at a tempo of 100 beats per minute. (**b**) The equivalent traces on the biomechanical drumming model from tracking experimental kinematics and dynamics from inverse kinematics (IK) and inverse dynamics (ID) solutions with a time-varying linear-quadratic regulator (TVLQR) controller. (**c**) The kinematics and dynamics resulting from a forward simulation performed with feedback from the TVLQR controller to maintain stability. The traces from the simulated solution produced nearly equivalent trajectories of the drumstick bead and joint centers with the largest discrepancies occurring at the elbow. The drumming model assumes a fixed shoulder joint center, whereas the real system exhibits slight translation resulting in small displacements of the elbow along the axial direction of the upper arm.

### 2.2 Modeling the Drumming Arm System

The equations of motion for a four degree of freedom drumming arm model (**Figure 1**) and their analytic Jacobians were derived in their explicit, continuous form using Kane’s equations of motion^[10]^ in Autolev^[11]^. The model consists of four revolute joints representing the sagittal plane rotational motion at the shoulder, *q*_*s*_, elbow, *q*_*e*_, wrist, *q*_*w*_, and fulcrum, *q*_*f*,_ connecting a chain of four rigid bodies representing the upper arm, forearm, hand, and drumstick—where the shoulder joint center is fixed in the Newtonian reference frame, N. Each generalized coordinate was actuated by a corresponding idealized torque actuator, *T*_*s*_, *T*_*e*_, *T*_*w*_, *T*_*f*_, while the contact interaction between the drumstick and the drumhead was modeled as a prescribed force, *F*_*d*_.

#### 2.2.1 Scaling the Model and Analyzing Experimental Techniques

Rigid body lengths were scaled with measurements of the drummer’s body segment lengths and rigid body masses were approximated by relating the measured total mass of the drummer to estimates of body segment masses via empirical relationships derived from water displacement measurements of 135 subjects^[12]^ and gamma-ray scans of 115 subjects^[13]^. Rigid body rotational inertias were estimated as uniformly distributed rods. A constrained inverse kinematics (IK) optimization was performed to find the shoulder, elbow, wrist, and fulcrum generalized coordinates, *q*, that minimized the error between the experimentally measured joint centers, *c*^*exp*^, of the drummer and the joint centers of the biomechanical model from forward kinematics, *c*:

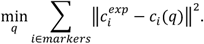

The generalized torques, τ, required to generate the kinematics from the IK solution were calculated via inverse dynamics (ID) by explicitly solving for τ in Kane’s equations of motion:

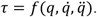

Finally, a coherent set of kinematics and dynamics representing the experimental drum stroke was acquired by tracking the estimated kinematics and dynamics from IK and ID simultaneously in a forward simulation. The forward simulation used a 4^th^-order Runge-Kutta discretization of the equations of motion^[14]^ with feedback from a discrete time-varying linear-quadratic regulator (TVLQR)^[15]^:

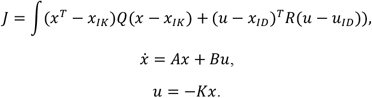

The state and control matrices, *Q* and *R*, of the TVLQR controller were tuned to emphasize precise matching of the kinematic state of the model from the forward simulation with the initial IK solution, and the *A* and *B* matrices of the linear dynamics model for 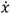 were chosen by linearizing the Jacobians of the model dynamics with respect to the IK, *x*_*IK*_, and ID, *u*_*ID*_, reference trajectories. The feedback gain matrix, *K*, was calculated using MATLAB’s (Mathworks, Inc.; Natick, MA) lqrd function for calculating the discrete-time feedback policy given continuous representations of the cost function and system dynamics. TVLQR was implemented because open-loop forward simulations with systems of rigid links are known to produce highly chaotic behavior, where numerical roundoff in the initialization and discretization causes deterministic systems to exhibit vastly different behavior over multiple simulations^[16]^.

### 2.3 Formulating the Trajectory Optimization Problem

The problem of finding the minimum effort drumming technique required to track a desired drumstick bead path was formulated as the following trajectory optimization problem, where the set of states, *x*, and control inputs, *u*, is identified which minimizes the effort of the drumming arm model while constraining the position of the bead of the drumstick to match a prescribed trajectory:

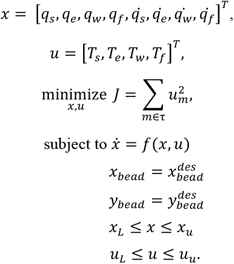

The state vector, *x*, of the model includes the four generalized coordinates and their first derivatives and the control vector, *u*, includes the four idealized torque actuators. Equality constraints were implemented to enforce the explicit dynamics of the model and to align the position of the bead of the drumstick with the prescribed trajectory. Inequality constraints were implemented to bound the states and controls to positions and motions that are feasible for the human arm and were selected using the TVLQR solution for the experimental drum stroke (**Figure 2**). The trajectory optimization problem was then transcribed into a nonlinear program (NLP) using Hermite-Simpson direct collocation^[17]^ and solved in IPOPT^[18]^.

### 2.4 Transcribing and Solving the Nonlinear Program

All computational procedures including data processing, optimization, and visualization were performed in MATLAB and all code is made publicly available on GitHub at: https://github.com/opearl-cmu/TrajOptDrums

Transcription of the trajectory optimization problem was performed using the publicly available OptimTraj toolbox^[9]^ which was edited to be directly interfaced with IPOPT using the mexIPOPT toolbox^[19]^. The result is a flexible library that supports a variety of transcription schemes as well as optimization with both MATLAB’s fmincon function and a state-of-the-art NLP solver (IPOPT) without leaving the MATLAB environment, which is particularly advantageous for debugging and tuning complex trajectory optimization problems.

## 3. Results

### 3.1 Experimental Kinematics and Dynamics

Tracking the experimental downstroke produces a coherent set of kinematics and dynamics that precisely characterizes the motion of the arm (**Figure 2**).

The stroke is first prepped and supported with larger body segments by actuating the shoulder and elbow before snapping the wrist and fulcrum down toward the drum and then breaking the downward motion of the forearm with the elbow.

### 3.2 Exploring Minimum Effort Alternatives

Solving for the minimum effort solution while tracking the prescribed path of the bead of the drumstick reveals a potential alternative to the experimentally performed technique (**Figure 3**) that possesses a lower effort requirement (**Figure 4**). Simultaneously tracking both the anterior-posterior (x) and vertical (y) paths of the bead resulted in a technique characterized by greater (mean difference ± standard deviation relative to the experimental data-driven TVLQR solution) shoulder flexion (+5.57 ± 3.16°), elbow extension (-9.17 ± 5.6°), and fulcrum flexion (+9.16 ± 5.90°); joint center trajectories that had larger ranges in the x-direction (+2.79 ± 1.00 cm); and actuation patterns that produced a lower root-mean-square (rms) cost (-14.83 N^2^m^2^).

**Figure 3.**
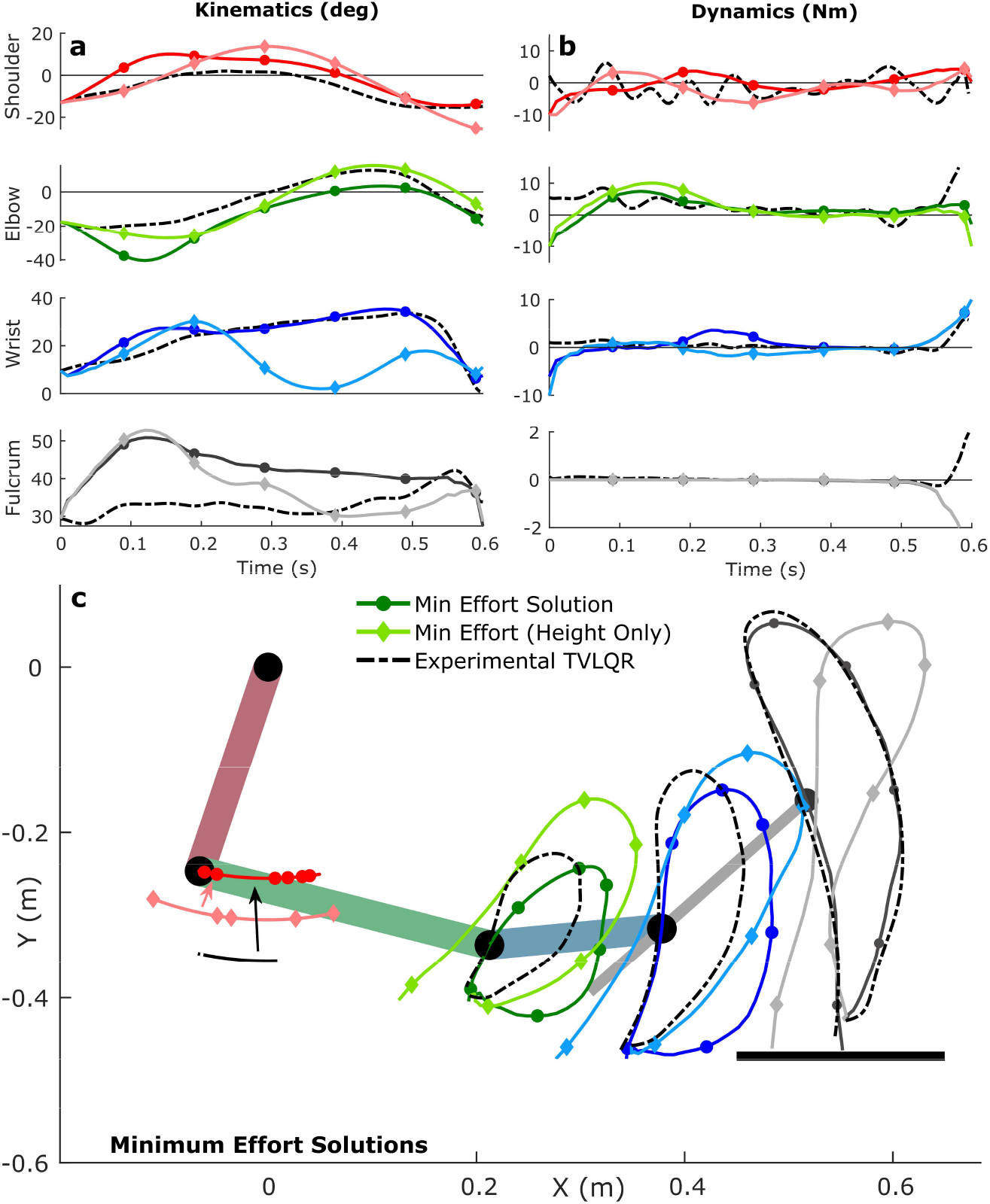
Minimum Effort Trajectory Optimization Solutions. A comparison of kinematics (**a**), dynamics (**b**), and trajectories of the drumming arm model (**c**) for the minimum effort solutions with (circles) and without (diamonds) tracking of the anterior-posterior bead trajectory against the experimental downstroke TVLQR solution (dashed lines). The minimum effort solution while tracking the full two-dimensional prescribed trajectory of the bead of the drumstick resulted in a stroke that is visually distinct from the experimental solution despite achieving an equivalent bead trajectory. The minimum effort solution while tracking only the height of the bead was even more visually distinct, as each joint center translated to the left of their initial starting positions.

**Figure 4.**
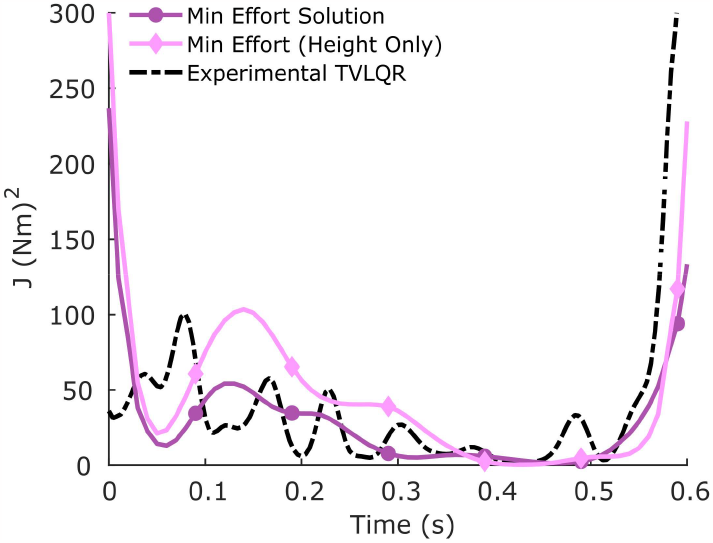
Minimum Effort Cost Comparison. The comparison of effort costs for each solution over the duration of the drum stroke shows that the minimum effort solution while tracking the bead’s full two-dimensional prescribed path resulted in a stroke with an overall lower effort than the experimental stroke. The effort also exhibited a less oscillatory pattern, indicating that the later part of the stroke more smoothly transitioned from using actuation at the joints to allowing gravity forces to propel the bead towards the drumhead. The minimum effort solution while tracking only the height of the bead resulted in a stroke with an overall higher effort requirement, exemplifying the possibility of discovering alternative locally optimal techniques that can be less efficient than even the experimentally recorded technique.

Tracking only the y-position of the bead allowed the optimizer to explore other local minima. The result was a technique characterized by joint center trajectories and a drumstick bead trajectory that translated in the negative x-direction (-6.59 ± 0.93 cm) relative to their starting point. The technique also required slightly higher RMS effort than the experimental solution (+8.24 N^2^m^2^) indicating that the optimizer converged to a local minimum that is worse than what it found when constrained to the full two-dimensional position of the bead.

### 3.3 Optimal Technique Across Body Type

A comparison of the minimum effort techniques for models scaled to the average male and female skeletal geometries and inertias shows visually distinct motion at the elbow and wrist joint centers resulting from key kinematic and dynamic deviations (**Figure 5**). The male technique has a much lower peak flexion at the elbow (-10.0°) and is generally more flexed at the shoulder (+1.17 ± 2.33°), wrists (+1.62 ± 2.24°), and fulcrum (+4.46 ± 3.68°). The overall pattern of the actuation is also unique as the resulting cost for the female stroke exhibits two peaks during the prep of the stroke whereas the male cost only exhibits a single large peak.

**Figure 5.**
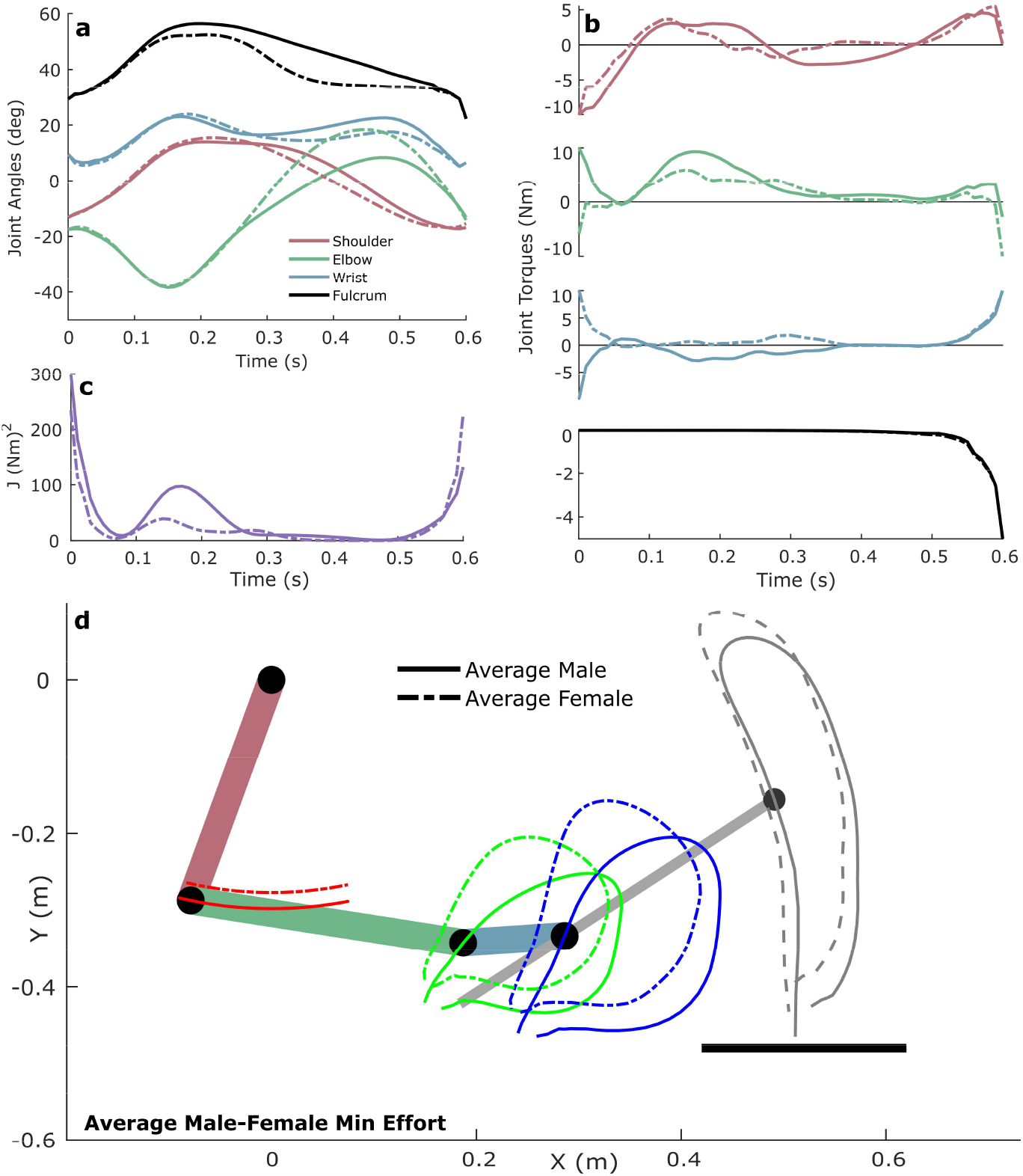
Optimal Technique Comparison Between Average Male and Female Body Types. The joint angles (**a**), joint torques (**b**), effort cost (**c**), and trajectories (**d**) of the minimum effort solutions using models scaled to match the skeletal geometry and inertia of the average male (solid lines) and female (dashed lines) body types. The entire two-dimensional position of the prescribed bead path was tracked, and the path was shifted for each body type so that the start of the path aligned with the starting position of the model’s bead given that both the male and female drumming arm models were initialized with the same state. The male and female minimum effort techniques exhibited distinct differences in the elbow and fulcrum angles as well as in the shape of the overall effort requirement (*J*). This suggests that, on average, males and females likely require slightly different drumming techniques to achieve maximum efficiency.

### 3.4 Optimal Technique Across Stroke Type

Simulations of single, double, and triple strokes provide a sense of the trends in minimum effort techniques across different stroke types (**Figure 6**). The function of each joint is mostly consistent between strokes, with the shoulder and elbow breaking the arm as it falls into the drum while the wrist and fulcrum pull the drumstick into the drumhead during contact. However, as more notes were played per stroke, less breaking (less positive torque) was done on average (with respect to the single stroke) at the shoulder (-0.21 Nm for the double stroke; -1.14 Nm for the triple stroke) and elbow (-0.49 Nm for the double stroke; -1.47 Nm for the triple stroke). The wrist also exhibited key differences between its optimal function for the single stroke and for the double and triple strokes. On average, the wrist helped break the system by providing a mostly positive torque during the double (0.45 Nm) and triple (0.05 Nm) strokes, whereas it provided a mostly negative torque (-0.64 Nm) to act in tandem with the fulcrum during the single stroke. The wrist angle also extended throughout the entirety of the double and triple strokes rather than flexing again after contact was made between the bead and the drumhead during the single stroke.

**Figure 6.**
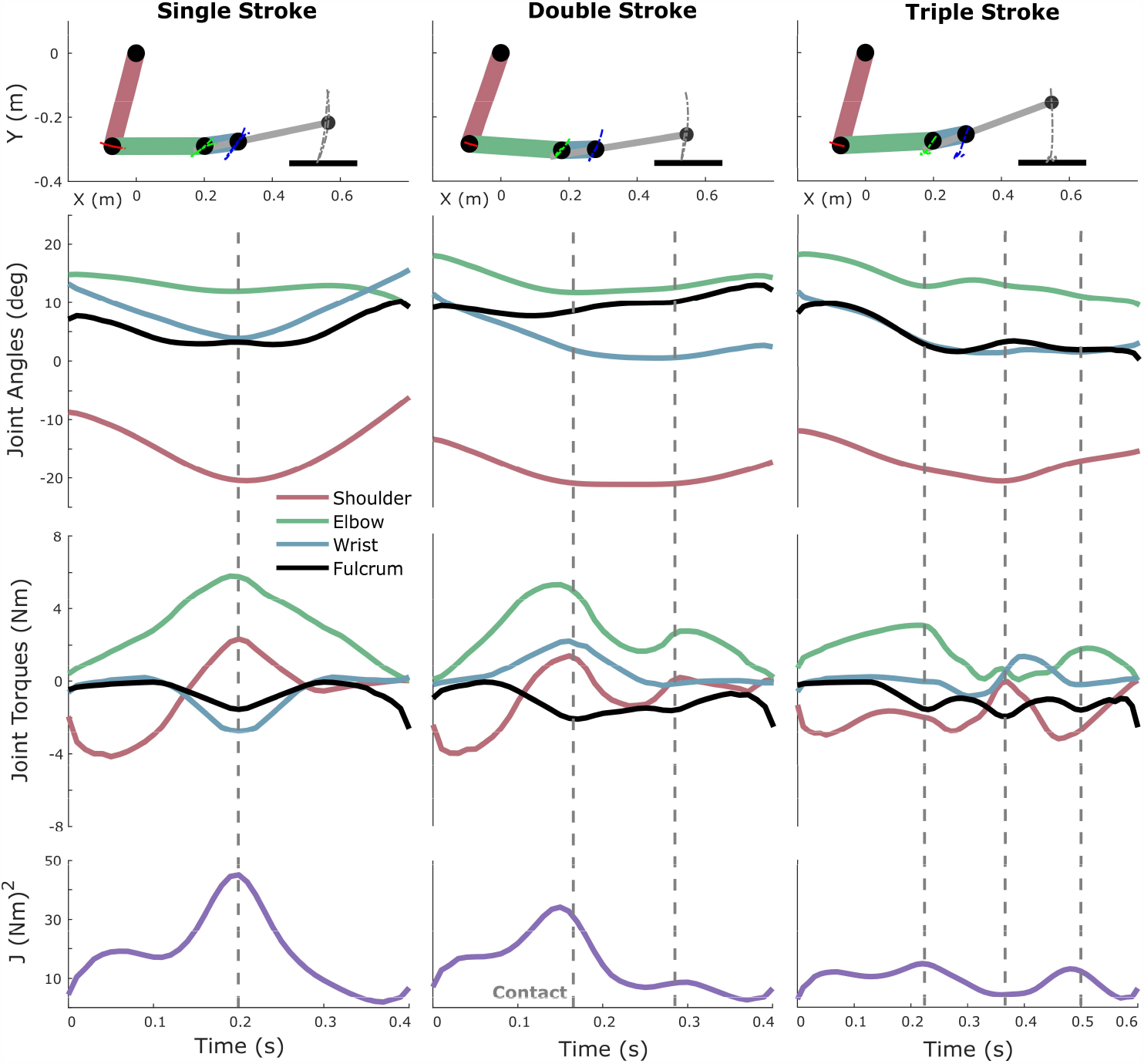
Comparison of Single, Double, and Triple Stroke Optimal Techniques. A comparison of the minimum effort solutions for a single stroke (left), double stroke (center), and triple stroke (right) reveals mostly consistent functions in the joints across very different motions. Each trajectory optimization was performed while only tracking the prescribed drumstick bead height, and the dashed vertical lines indicate where impact with the drum occurred. As more notes were played, the shoulder and elbow performed less breaking. As the arm switched from performing a single stroke to a series of strokes, the wrist switched from working synchronously with the fulcrum to working against it. This suggests that general trends in the optimal function of each drumming lever do exist, but that the optimal function for each lever can switch depending on the type of stroke being performed.

## 4. Discussion and Implication

### 4.1 Significance and Impact

Having the capability to analyze a drummer’s technique for areas of suboptimality quickly and efficiently could revolutionize how percussion develops and continues to shape the world. In this study, I have demonstrated how trajectory optimization with biomechanical models can be used to analyze the efficiency of experimental drum strokes, search for new and improved techniques, and compare optimal techniques across different body types and strokes. I have also made available a user-friendly MATLAB library through which others can easily perform a quantitative analysis of drumming technique by prescribing the path for the bead of the drumstick, setting the parameters of the biomechanical model, and adjusting the trajectory optimization problem to meet their needs. The library then automatically transcribes the problem into an NLP using either trapezoidal, Hermite-Simpson, or Runge-Kutta direct collocation transcription methods and solves the NLP with fmincon or IPOPT.

### 4.2 Limitations

The study was limited to a simplified drumming arm model with a low-dimensional state space. Precisely capturing the real-world behavior of the arm would likely require a more comprehensive model, as analogous analyses of human gait for clinical applications often require high degree of freedom representations of the legs with subject specific inertial parameters and musculotendon actuators^[20–24]^. However, employing a simple drumming arm model to capture the core behaviors of the system is useful because it requires lighter code, is more computationally efficient, and enables faster iteration. This is advantageous for quickly extracting the general governing principles of the real-world drumming arm system while avoiding overfitting^[25,26]^. Yet, a similar analysis to the one performed in this study could potentially be performed in a more precise, high-dimensional space by combining more representative models of the arm, such as those developed by the OpenSim community^[27,28]^, with libraries specially designed for solving trajectory optimization problems with high-dimensional models incorporating multiple musculotendon actuators and complex contact force interactions^[29,30]^.

### 4.3 Identifying Better Drumming Approaches

Comparing the experimental drum stroke with the solution to the minimum effort trajectory optimization problem via the reduced-order model revealed that the experimental drum stroke used 30% more effort than the optimal solution. This difference in effort along with the stylistic shift in the technique stemming from kinematic differences occurring at the shoulder, elbow, and fulcrum highlights the advantage of using trajectory optimization to analyze drumming technique and search for more efficient alternatives. A percussion instructor could harness such information to provide specific feedback to a student to help them adjust and hone their technique. For the stroke analyzed in this study, it would be appropriate to instruct the drummer to “use less elbow” (more extension) and instead “use more shoulder” (more flexion) to drive the stroke while “relaxing their fulcrum” (allowing more flexion) during the prep of the stroke, which together would enable the arm to more efficiently leverage gravity forces to redirect and drive the stick to the drum.

Exploration of alternative techniques must be done with care because of the nonlinearity of the system dynamics. Performing the same minimization without constraining the bead of the drumstick to match the experimental path in the anterior-posterior direction resulted in a technique that was stylistically unique but required 13% more effort than the experimental technique. This exemplifies the possibility that the optimizer can converge to locally optimal techniques that may not be better than the experimentally recorded technique, depending on the constraints, bounds, and initial guess provided to the NLP solver. A large drawback of performing trajectory optimization on systems with nonconvex equations of motion is that there is no way to guarantee that a particular solution is the global optimum^[31]^. However, extensive probing of the cost function through repeated iterations or an additional global pattern search can help one better evaluate the solution space and find a more appropriate local minimum^[32,33]^.

### 4.4 Exploring Technique Accessibility

The optimal techniques for models scaled to match the average male and female body types deviated significantly at the elbow and fulcrum. Access to such information is helpful for understanding whether differences in the optimal techniques of males and females is a contributing barrier to entry for females in a field that is traditionally male dominated^[34]^. When drumlines or percussion sections treat the optimal male technique as the benchmark for proper form, it is likely that they are implicitly forcing female members and auditionees to adapt to a technique that is not as efficient for their body type. An important trade-off revealed in this study is the need for greater flexion at the elbow for the average female than for the average male given the same bead trajectory criteria. This greater flexion angle suggests that the elbow will be more rigorously loaded over time for female players than male players if they are expected to move the stick in exact same way. This additional loading requirement over thousands of repetitions could have significant ergonomic consequences and should be considered when instructing or assessing performance in order to avoid unfairly favoring one body type over another.

### 4.5 Designing Coherent Drumming Strategies

Observing the optimal techniques across a single, double, and triple stroke shows that each joint has a mostly consistent optimal function, even across motions with unique dynamics requirements. This bolsters the argument for teaching percussion through a top-down approach that adopts a common set of principles which the individual percussionists then adapts to meet the specific performance demands of a particular stroke or rudiment^[3]^, rather than teaching the technique for each stroke or rudiment in isolation. However, the noteworthy switch in the wrist’s optimal function from actuating synchronously with the fulcrum during the single stroke to working opposite the fulcrum during the double and triple stroke to help break the motion at the base of the drumstick suggests that it is also important to incorporate slight, yet targeted changes in technique between different stroke types to minimize effort. A traditional top-down approach to teaching percussion might not be capable of capturing these subtle differences in the optimal technique across different stroke types, but trajectory optimization is well suited for identifying these differences and could therefore serve to augment top-down paradigms by quantitatively informing more subtle areas of improvement.

### 4.6 Conclusion

This work illustrates both the power of the qualitative optimization of drumming technique that has taken place over the past seven thousand years and the power of using quantitative optimization to validate, inform, and reshape existing drumming paradigms. Intuition developed qualitatively can be used to inform and bound biomechanical simulations, which can then serve to help iterate on and improve human intuition. If adopted by the percussion community, quantitative consultations using biomechanical models and trajectory optimization could assist percussionists in their endeavor to perform at the highest level while maximizing efficiency and preserving long-term health.

## Acknowledgments

I would like to thank Carnegie Mellon University’s National Biomechanics Day for inspiration to think outside the box.

## Data & Code Availability Statement

All code, experimental data, and simulated results associated with this publication are publicly available through GitHub at https://github.com/opearl-cmu/TrajOptDrums.

